# Multimodal imaging of human fetal brain development at the mesoscopic scale using 11.7 T ex vivo MRI

**DOI:** 10.1101/2025.09.08.669657

**Authors:** Lucas Arcamone, Cyril Poupon, Homa Adle-Biassette, Suonavy Khung-Savatovsky, Marianne Alison, Jessica Dubois, Lucie Hertz-Pannier, Yann Leprince

## Abstract

We present the first release of p-HCP (Prenatal Human Connectome Patterns), an imaging dataset of human fetal brain development covering the second half of gestation. This dataset was acquired ex vivo using magnetic resonance imaging (MRI) at ultra high field (11.7 teslas), and includes whole-hemisphere T_2_-weighted images at 100 µm isotropic resolution, quantitative relaxometry (T_1_, T_2_, and 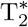), and high angular resolution diffusion-weighted images for multiple b-values at 200 µm. Brains larger than the workspace of the small-bore scanner were sectioned into blocks, acquired blockwise, and digitally reconstructed using a dedicated semi-automatic method.

This initial data release includes three gestational ages (18, 27, and 31 post-conceptional weeks) with a complete set of anatomical images, relaxometry maps, and diffusion-based microstructure measurements. This dataset offers new opportunities to investigate neurodevelopmental processes that have not yet been explored with full three-dimensional coverage at this resolution by MRI, and may serve as a multimodal mesoscopic reference template for the fetal brain.

## Background & Summary

The second half of human gestation, corresponding approximately to 18 to 40 post-conceptional weeks^1^ (PCW) is a critical period for fetal brain development. This period is marked by the emergence of transient fetal compartments that support key neurodevelopmental processes such as neuronal migration, axonal growth, synaptogenesis, and gliogenesis [1–4]. These dynamic events lay the foundation for the structural and functional architecture of the postnatal brain [5–7].

Our current understanding of these cellular and molecular processes is derived predominantly from histological analyses, particularly the microscopic examination of stained tissue sections [2, 8–11]. These techniques provide high-resolution two-dimensional insights into tissue microstructure and cytoarchitecture. However, their spatial coverage is inherently limited, and densely sampled, whole-brain three-dimensional histological datasets are still lacking for the human fetal brain. Recent advances in image processing have enabled the reconstruction of three-dimensional microscopic datasets from sparsely sampled histological sections, using high-resolution magnetic resonance imaging (MRI) as a structural reference framework [12]. These multimodal approaches hold great promise for bridging the gap between micro- and macro-scale observations. However, their application to the human fetal brain remains extremely difficult, primarily due to the rarity of well-preserved fetal specimens, the need for highly specialized and interdisciplinary expertise, and the considerable financial and logistical costs associated with such large-scale integrative studies. The absence of such integrated datasets represents a major limitation for developmental neuroscience, as it hinders the creation of comprehensive spatiotemporal atlases of the human fetal brain. These resources are essential for correlating molecular, cellular, and structural events across developmental stages, and for informing both experimental models and clinical interpretations. Bridging this gap will require not only methodological innovation, but also coordinated efforts to access, process, and standardize rare fetal brain samples.

In parallel, in utero fetal MRI has experienced rapid advances in recent years [13–17], driven by the development of ultra-fast sequences and sophisticated reconstruction methods that yield full-brain three-dimensional (3D) datasets, including morphological, diffusion-weighted, and even functional imaging. However, in utero fetal MRI remains constrained to a macroscopic resolution due to several limiting factors, including spontaneous fetal movements, specific absorption rate (SAR) limits, and limited acquisition time.

Ex vivo MRI at ultra-high field is a compelling alternative for achieving higher resolutions, as it is not limited by SAR nor acquisition time [18]. Prolonged scanning allows the acquisition of high quality 3D multimodal data at a mesoscopic scale (≈ 100–200 µm), bridging the gap between macroscopic in vivo imaging and microscopic histological analysis. Such mesoscopic data pave the way for a deeper understanding of the relationships between MRI signal characteristics and underlying biological parameters, albeit within the inherent limitations of comparing in vivo and ex vivo datasets. Recently, studies have used MR mesoscopic data to explore correlations between histological data and quantitative MRI by creating biophysical models, opening a new field called MR histology [19, 20]. Notably, effective transverse relaxation time 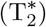 has been shown to correlate strongly with both iron concentration and myelin volume fraction [21, 22]. Longitudinal relaxation time (T_1_) is sensitive to the restricted mobility of water molecules and chemical exchange, making it a useful marker for macro-molecule concentration, such as lipids that constitute the myelin sheath [23]. In this context, multiparametric approaches have shown promising results in characterizing tissue-specific signatures in adults and in infants [24–26], but to the best of our knowledge such methods have not yet been applied to fetal brains. Similarly, diffusion imaging also provides information on the tissue microarchitecture, and can be used to explore both the cytoarchitecture of the brain as well as white matter organization [27, 28]. Notably, diffusion tensor imaging (DTI) metrics are known to strongly correlate with fibre density and orientation distribution [29, 30]. More recent models allow a finer characterization of the microstructure [31–33].

Due to the tissue fixation and lower temperature at which post-mortem imaging is performed, water diffusivity is markedly reduced, necessitating the use of high b-values to explore fine details of the tissue microstructure. Such high b-values require strong magnetic field gradients above 700 mT/m, which are currently only achievable with small-bore MRI scanners designed for rodents. As a result, the few existing studies of fetal brains with ultra-high-field MRI have been limited to small brains, generally under 20 PCW, or have focused on isolated brain structures [34]. This restriction stems from the fact that more advanced fetal brains are physically too large to fit within these scanners. In the present project, we overcome this limitation through a novel methodology, adapted from the Chenonceau project, originally developed for the adult human brain [35, 36]. Our approach involves the blockwise acquisition of a dissected specimen, followed by in silico reconstruction of the whole-brain image.

Using this methodology, the present study introduces an unprecedented developmental series of multimodal mesoscopic anatomical and quantitative MR datasets of ex vivo human fetal brains, covering the late second and third trimesters of gestation. This first data release focuses on three ages: 18, 27, and 31 PCW. It includes T_2_-weighted images and quantitative 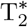 maps (100 µm 3D isotropic) of the whole samples, as well as quantitative T_1_ maps and T_2_ maps, and diffusion microstructure maps (200 µm 3D isotropic). Future data releases will incorporate tractography-based reconstructions of white matter pathways, advanced multi-shell diffusion models, histological data acquired from the same specimens, and additional developmental stages. This dataset, illustrated in Figure 1, provides a unique opportunity to characterize brain structures and their development throughout gestation with unprecedented detail at the whole-brain level, particularly transient fetal compartments involved in neurogenesis, neuronal migration and axonal growth. Quantitative multimodality enables microstructural modelling, paving the way for a better understanding of the brain tissue composition and neurodevelopmental dynamics during the last two trimesters of gestation (Figure 2). The combination of isotropic mesoscopic resolution, multimodal contrasts, and whole-hemispheric coverage makes this dataset a strong candidate to serve as a reference atlas for the study of late fetal brain development.

**Figure 1:**
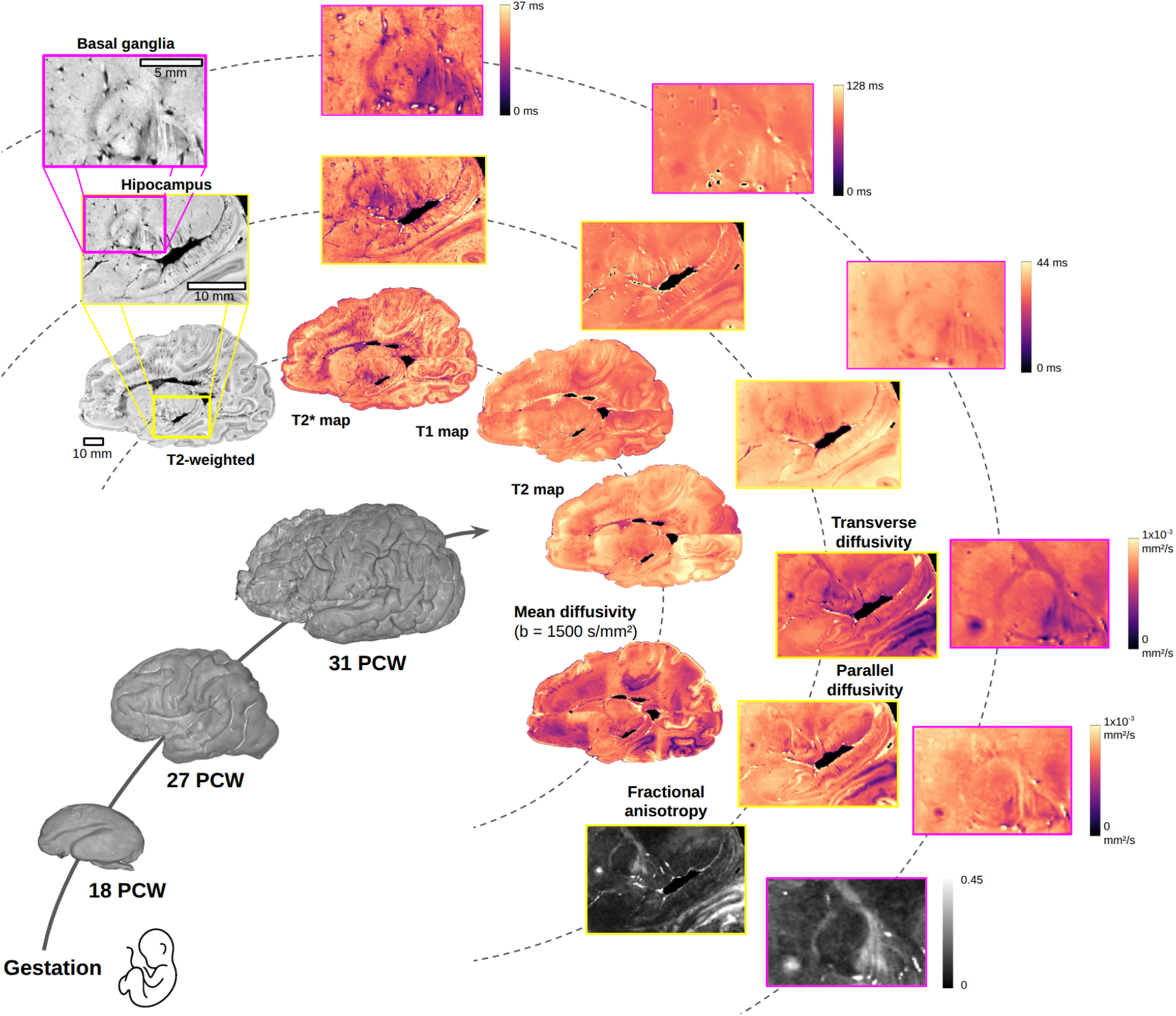
Overview of available modalities and specimens, with a focus on the hippocampus and the basal ganglia in the 31-PCW specimen. The figure shows the outer brain surface of the three specimens with a volume rendering of the T_2_-weighted images, and a choice of modalities for the 31-PCW specimen. Inner ring: a whole sagittal slice; intermediate ring (yellow frames): highlights a zoomed-in view of the basal ganglia and the hippocampus; outer ring (purple frames): provides a further magnified view focusing specifically on the basal ganglia. PCW = post-conceptional weeks.

**Figure 2:**
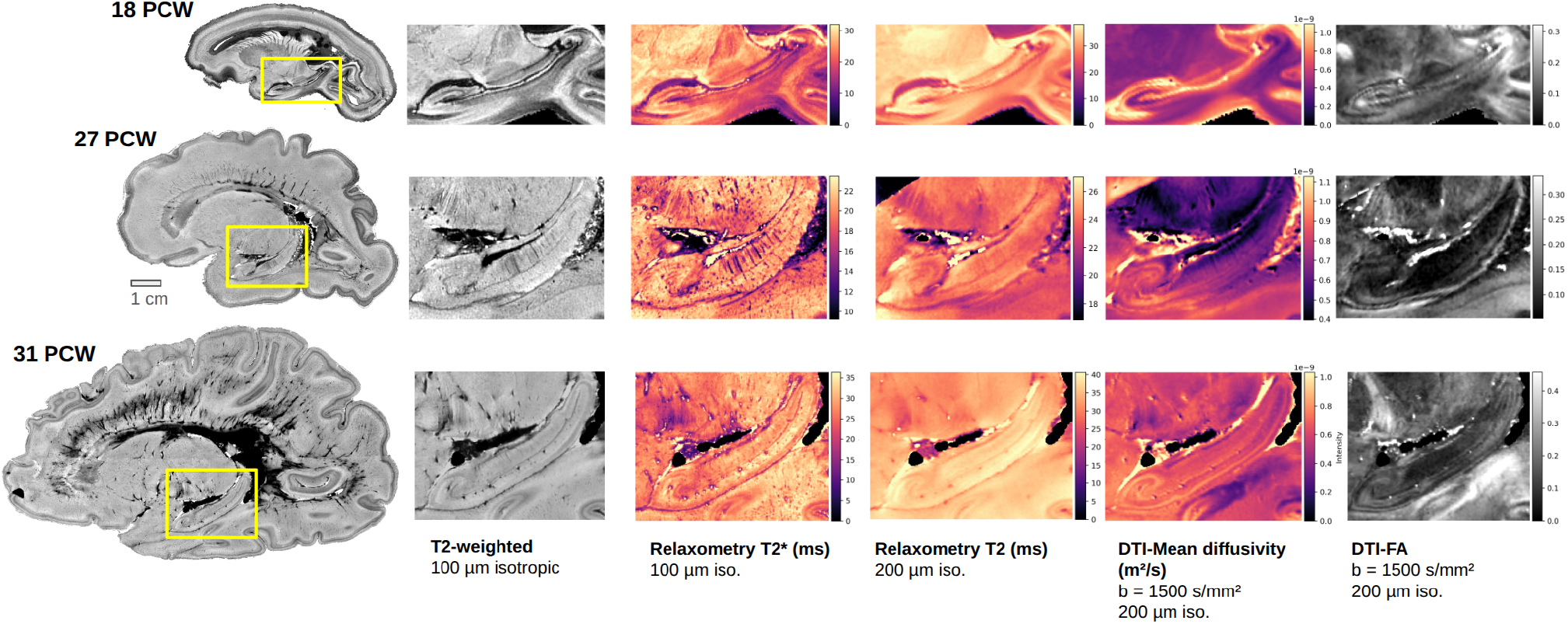
Three stages of brain development: focus on the hippocampus. Left column: whole-hemisphere – T_2_-weighted sagittal slices. Other columns: zoom on the hippocampus on several modalities. DTI = diffusion tensor imaging, FA = fractional anisotropy.

## Methods

Figure 3 provides an overview of the acquisition and reconstruction workflow. Each brain was first imaged in situ using 3 T MRI, within few hours after death, as part of a routine virtopsy protocol. The brain was then extracted, fixed in formalin, and the hemispheres were separated. After the fixation period, one hemisphere was rinsed, doped with a gadolinium-based contrast agent, and imaged with 7 T MRI to obtain a submillimetric whole-hemisphere reference image. The hemisphere was subsequently sectioned into smaller blocks to fit within the small bore of an 11.7 T MRI scanner. Each block was imaged across multiple fields of view (FOVs). Finally, the whole-hemisphere volumes were reconstructed by stitching all the FOVs, using a three-step semi-automatic registration pipeline. Each of these processing steps is detailed in the following subsections.

**Figure 3:**
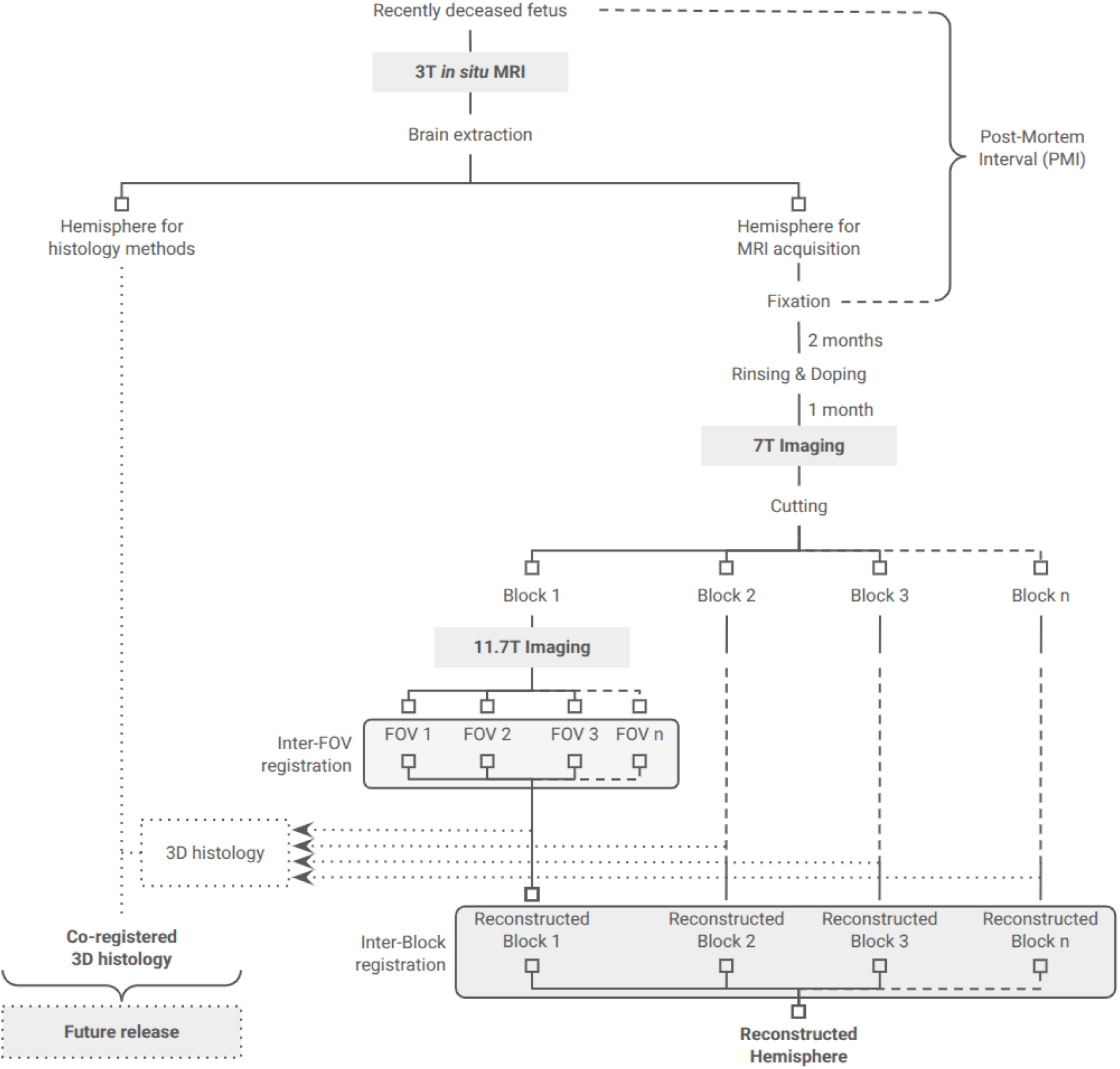
Overview of the processing, acquisition, and reconstruction pipeline. FOV = field of view.

### Specimens

This study was conducted in accordance with the principles outlined in the Declaration of Helsinki. Informed consent was granted by the parents, and the handling of human brain tissue was declared to the competent regulatory authority (CODECOH declaration DC-2022-5118).

The specimens were selected by the department of Fetal Pathology at Robert-Debré hospital (Paris, France) from normotrophic fetuses following miscar-riage, medical termination of pregnancy, stillbirth, or neonatal death within the hospital. Exclusion criteria included any condition—genetic or acquired—known to significantly impact brain development, as well as any visible lesions or malformations visible on the in situ MRI.

This first data release includes three specimens representing key stages of the fetal brain development (18, 27, and 31 PCW). The characteristics of these specimens are summarized in Table 1.

**Table 1:** General characteristics of the specimens.

| Identifier | Post-con-<br>ceptual<br>age at death | Sex | Cause of death | Observations | Vascular<br>rinsing | En-<br>cephalic<br>weight | Post-mortem<br>interval | Imaging protocol |
| --- | --- | --- | --- | --- | --- | --- | --- | --- |
| sub-t18a | 18.1 w | F | Miscarriage<br>on acute<br>chorioamni-<br>otitis |  | yes | 40 g | 36 h | whole brain<sup>a</sup><br>1 block, 1 FOV<br>150 h acq. time<sup>c</sup> |
| sub-t27arh | 27.0 w | F | Medical ter-<br>mination of<br>pregnancy | 22q11 dele-<br>tion<sup>b</sup>, cardiac<br>malformation | no | 229 g | 25 h | right hemisphere<br>2 blocks, 4 FOV<br>600 h acq. time<sup>c</sup> |
| sub-t31alh | 30.9 w | M | *Placenta*<br>*abruptio* | Chronic<br>hypoxia | yes | 319 g | 31–43 h | left hemisphere<br>2 blocks, 6 FOV<br>1000 h acq. time<sup>c</sup> |

<sup>a</sup>The hemispheres of sub-t18a were not separated, because the entire brain could fit into the 11.7 T scanner; also 7 T and 3 T in situ imaging are missing for that specimen.
<sup>b</sup>despite the genetic anomaly and given the rarity of fetal death at this age, we have included this brain as a pseudo-typical specimen, on the basis that only mild structural brain anomalies have been described for 22q11 brains having normal brain size [37] (here brain weight is between the 50–70th percentile).
<sup>c</sup>Total acquisition time for all FOVs at 11.7 teslas.

### Sample preparation and handling

#### Extraction, fixation & doping

After in situ 3 T imaging (described below), two of the specimens underwent vascular rinsing (see Table 1) via intracarotid perfusion with phosphate-buffered saline (PBS), continued until the fluid returned through the brachiocephalic vein. This procedure, commonly used in adult autopsy protocols, aims to reduce MRI artefacts caused by residual blood. However, it was subsequently omit-ted for later specimens due to technical difficulties in fetal cases and concerns about potential damage to the fragile tissue. Following extraction from the skull, the brains were immersion-fixed in 12% buffered formalin. In two cases, one hemisphere was dedicated to histological analyses (not included in the present data release), while the other was processed for MRI. In the third case, the entire brain was scanned before histological processing. A minimum of two months of fixation at ambient temperature was required before imaging. After fixation, the tissue was rinsed and doped by immersion in a 0.1 mol/L PBS solution supplemented with 2 mmol/L gadolinium chelate (Gd-DOTA) and 0.01% (w/w) sodium azide, and stored for at least one additional month at 5°C. During MRI acquisitions at both 7 T and 11.7 T, the tissue was immersed in fluoropolymer oil (Fluorinert™ FC-40), which minimizes magnetic susceptibility artefacts while generating no MR signal [38].

#### Cutting & Storage

The method of cutting and storing the hemispheres was developed to limit handling as much as possible, to minimize damage to these particularly fragile samples, which were susceptible to tearing under their own weight. We acquired the reference image at 7 T (see below) to guide the definition of cutting plane(s), which were selected to minimize the number of necessary FOVs while preserving at best the integrity of brain structures of interest. A custom cutting die (see Figure 4.e) was designed using computer-aided design software (FreeCAD), and fabricated in photopolymer resin using a stereolithography (SLA) 3D printer. This allowed us to cut hemispheres precisely along the chosen cutting planes. The resulting blocks were stored and imaged in a custom SLA-printed container consisting of a sealed cylindrical outer container, and upper and lower inner holders (see Figure 4.b and .c). The outer container was a simple 50 mm inner diameter cylinder, with a sealed lid (see Figure 4.a) clamping onto the motorized bed of the MRI scanner (see Figure 4.d), enabling total immersion of the block in fluoropolymer oil and precise sample positioning. The lower inner holder was a generic semi-circular part used to hold gauze against the tissue and prevent block from rotating. Due to the high density of the fluoropolymer oil (1.9 g/ml), buoyant forces pushed the tissue strongly against the upper inner holder, which was shaped to closely fit each block’s geometry. Each block remained in this container until all acquisitions had been completed and quality-checked, permitting precise repositioning and limiting the risk of mechanical damage from repeated handling. Containers were stored at 5°C but were brought to room temperature for 24 hours prior to each imaging session, to allow the tissue to reach thermal equilibrium.

**Figure 4:**
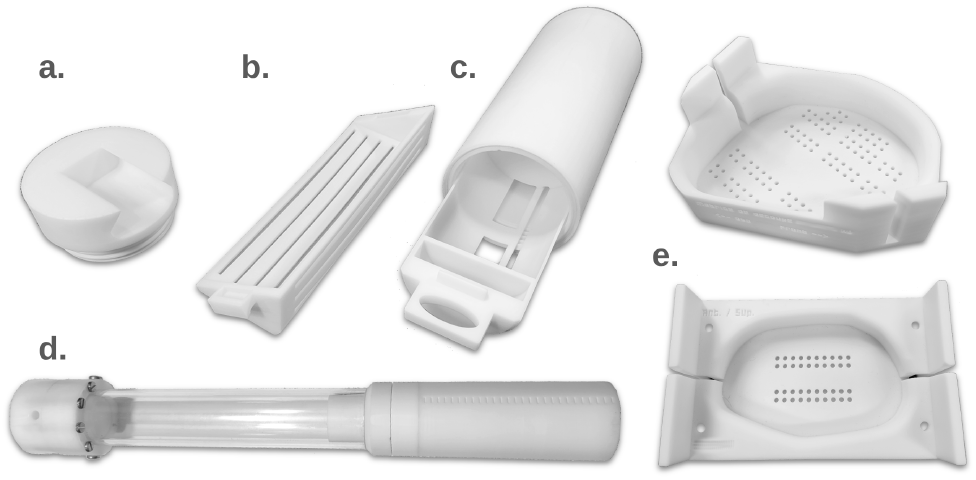
Custom 3D-printed hardware. a. Sealing cap of the external container, b. Upper inner holder, c. External container with lower inner holder, d. Container attached to the MRI motorized bed, e. Cutting die for sub-t27arh (top) and sub-t31alh (bottom). SLA = stereolithography.

### MRI data acquisition

#### 3-tesla in situ imaging

As part of a routine virtopsy protocol, bodies were imaged using a 3 T clinical MRI scanner (Ingenia, Philips Medical System, Best, The Netherlands) at Robert-Debré hospital. A T_2_-weighted 3D driven equilibrium (DRIVE) fast spin echo (FSE) sequence was used (resolution ≈ 0.6 mm). These images were used to select the specimens to be included in the study.

#### 7-tesla whole-hemisphere imaging

Whole-hemisphere reference images were acquired on a 7 T clinical MRI scanner (Magnetom 7-tesla investigational device, Siemens Healthineers, Erlangen, Germany) at the NeuroSpin neuroimaging centre, with a 3D T_2_-weighted turbo spin echo (TSE) sequence (matrix size 512 *×* 512 *×* 230, resolution 400 to 500 µm depending on the FOV, TR = 500 ms, TE = 45 ms, echo train length = 7, 4 repetitions, acquisition time = 28 min). The FOV was adjusted to the size of each specimen. 7 T MRI was not performed on small specimens not needing to be cut (presented in this first release: sub-t18a).

#### 11.7-tesla blockwise imaging

Mesoscopic scale acquisitions were performed on an 11.7 T small-bore MRI (BioSpec 117/16 USR Bruker MRI, Ettlingen, Germany), whose gradient system can reach 760 mT/m. A 60-mm inner diameter transmit/receive volumetric quadrature coil was used, ensuring good B_1_ homogeneity, a high signal in deep structures, and wide coverage for large regions of interest. ParaVision 6.0.1 was used to perform the acquisition.

Even with a wide coverage, a single block of tissue could require several FOVs because of the limited space available inside the coil (60 mm along the tunnel direction). Adjacent FOVs were overlapped by at least 15 mm to facilitate reconstruction. The total acquisition time at 11.7 T is described in Table 1.

T_2_-weighted high-resolution imaging (100 µm isotropic) was performed with a 3D spin echo sequence (TE = 55 ms, TR = 350 ms, bandwidth = 179 Hz, flip angle = 90°, acq. time = 21 h).

T_1_ relaxometry (200 µm isotropic) was performed with the Variable Flip Angle Spoiled Gradient Echo method (VFA-SPGR), using a series of 70 3D Fast Low Angle Shot (FLASH) sequences (TE = 4.03 ms, TR = 10 ms, bandwidth = 370 Hz, flip angle = [20–90]° (70 values), total acq. time = 13 h), combined with a map of transmit B_1_ field acquired with the 3D Actual Flip Angle (AFI) method (TE = 3.3 ms, TR_1_ = 15 ms, TR_2_ = 45 ms, isotropic voxel size = 600 µm, bandwidth = 538 Hz, flip angle = 60°, acq. time = 6 min).

T_2_ relaxometry (200 µm isotropic) was performed using a 3D multi-echo spin echo sequence (TE = [10–200] ms (20 echoes), TR = 500 ms, bandwidth = 370 Hz, flip angle = 90°, acq. time = 7 h 20 min).

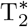 relaxometry (100 µm isotropic) was performed using a 3D multi-echo gradient echo sequence (TE = [5.2–81.7] ms (10 echoes), TR = 100 ms, bandwidth = 179 Hz, flip angle = 50°, acq. time = 6 h). To reduce signal loss due to susceptibility artefacts, particularly around air bubbles, we improved the spatial resolution to 100 µm, which is finer than the resolution of other quantitative scans.

For diffusion imaging (200 µm isotropic), High Angular Resolution Diffusion Imaging (HARDI) data were acquired on 3 shells using 3D Pulsed Gradient Spin Echo (PGSE) sequences with a segmented 3D Echo Planar Imaging (EPI) reader module (TR = 250 ms, TE = 24.05 ms, bandwidth = 1071 Hz, number of segments = 30, flip angle = 90°, b = 1500 / 4500 / 8000 s/mm^2^, with 25 / 60 / 90 diffusion directions respectively, total acq. time = 100 h).

### Parametric mapping

The quantitative maps were reconstructed in the native space of acquisition for each FOV. Processing scripts are publicly available, see the Code Availability section.

#### T_1_ mapping

As a pre-processing step of T_1_ mapping, the 70 VFA-SPGR images acquired with varying flip angles were affinely registered to the first one, to compensate for progressive tissue flattening during this 13-hour long acquisition. The T_1_ maps were generated using the VFA method with 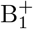 map correction, with maps of 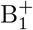 reconstructed from the AFI sequence [39].

As we systematically observed lower T_1_ values in tissue regions at the top of each FOV (further discussed in Technical Validation), we estimated a smooth multiplicative bias field using the N4 algorithm [40], isolated affected regions by fitting a Gaussian mixture model to the bias field, and selectively applied a multiplicative correction to those regions of low estimated T_1_.

#### T_2_ & 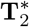 mapping

A mono-exponential fit was used for both quantitative T_2_ and 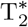 maps. The first echo was excluded from the T_2_ fit [41]. No registration was necessary, as all echoes were acquired simultaneously in multi-echo sequences.

#### Diffusion imaging

The diffusion-weighted images were pre-processed to correct for eddy current artefacts and motion (slow flattening of the tissue during the very long acquisitions) using affine registration. Non-local means filtering was applied to reduce the noise, using a Rician noise model, the noise variance being estimated from the mean square of the signal in the background [42]. We further bias-corrected the diffusion-unweighted (b = 0) images to compensate for a systematic localized signal loss—further discussed under Technical Validation. This was done by using N4 [40] to estimate a bias field, isolating affected regions by fitting a Gaussian mixture model to the bias field, and selectively applying a multiplicative correction to those regions of signal loss. For each b-value, a tensor model was fitted [43] and several parameters were extracted: Mean diffusivity (MD), Fractional Anisotropy (FA), Parallel Diffusivity and Transverse Diffusivity (also known as axial and radial diffusivity, respectively). For each b-value, the Analytical Q-Ball Imaging model [44] was also fitted, and Generalized Fractional Anisotropy (GFA) was computed.

### Image registration

In this work, the crucial step was the ability to recompose a whole-brain image from multiple partial FOVs of multiple tissue blocks. This was realized by first estimating the spatial transformation from each FOV (“initial space”) to a common anatomical space (“final space”) using image registration techniques, then merging the information from all FOVs by an adapted image fusion strategy.

To estimate the spatial transformation that brings images from the initial space to the final space, we developed a semi-automatic methodology based on linear and non-linear registrations (ANTs SyN symmetric diffeomorphic registration [45], RRID:SCR_004757). The method comprises three main steps, summarized in Figure 3: firstly, we realigned the data onto a reference modality in the initial space (intra-FOV registration), then we reconstructed each tissue block from the overlapping FOVs (inter-FOV registration), and we finally developed an original method to reconstruct the entire hemisphere from the reconstructed blocks (inter-block registration). This last step was the most challenging, as the blocks had no overlapping areas.

#### Intra-FOV registration

Within each FOV, we aligned all modalities to the 11.7 T high-resolution T_2_-weighted image, chosen as reference because of its high contrast and consistent quality. Indeed, while the blocks were firmly maintained by the container, we still observed a subtle progressive tissue deformation along the vertical axis over the duration of a complete acquisition session (150 hours per FOV). To compensate for these intra-FOV geometry variations, we applied affine registration between each modality and the T_2_-weighted reference, using a Mutual Information cost function to handle this cross-modal registration. This step also allowed us to focus only on the T_2_-weighted reference images in subsequent registration steps, improving their robustness and accuracy thanks to the use of monomodal registration. This intra-FOV registration step was also essential to compensate for bulk displacements in a few cases where a specific FOV had to be acquired over several non-consecutive sessions, either for organizational reasons or due to failed acquisitions.

#### Inter-FOV registration

This registration step involved merging adjacent overlapping FOVs to reconstruct full-block images (see Figure 5). Inter-FOV registration allowed for correction of the uncertainty of the translation made by the MRI bed, and the flattening differences and subtle non-linear deformations that appeared between acquisition sessions.

**Figure 5:**
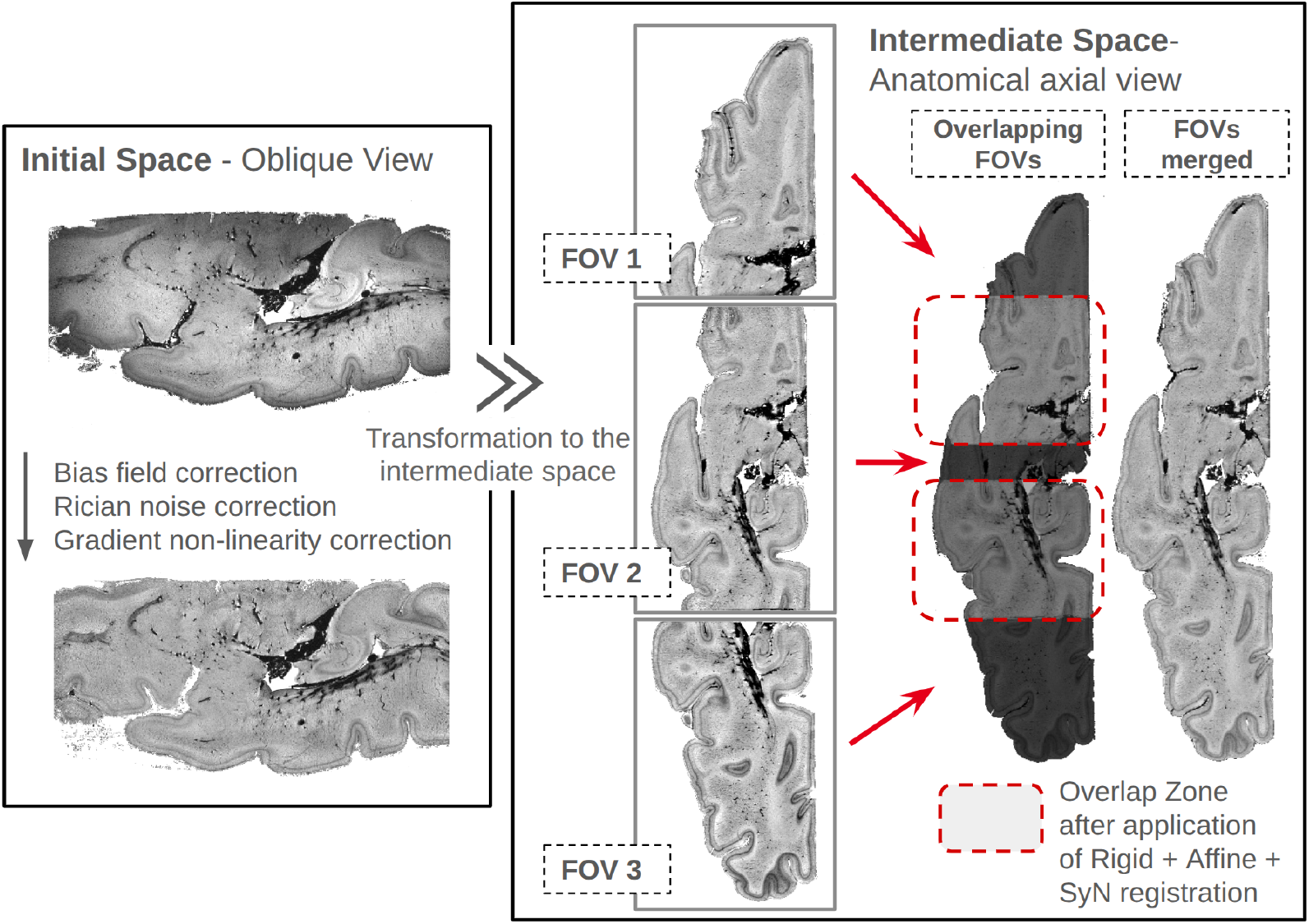
Schematic of inter-FOV registration. Illustrated with the inferior block of sub-t31alh. Left frame: image before and corrections in the initial space. Right frame: registration process to reconstruct the block from 3 FOVs in the intermediate space. FOV = field of view.

To maximize the quality and reproducibility of the registration steps, we first cleaned data for noise, bias, and geometric distortions due to non-linearity of the gradients. Non-local means filtering was used to denoise the data, using a Rician noise model. N4 was used to perform bias field correction [40]. Gradient non-linearity correction was performed by generating an ANTs deformation field based on the correction method implemented in ParaVision 7. This approach preserved format consistency throughout the reconstruction pipeline, facilitating the later composition of non-linear deformation fields.

For each FOV doublet sharing an overlap space, we isolated the overlap area with a mask, whose voxels were exclusively taken into account when optimizing the registration metric. In addition, the physical translation performed with the preclinical MRI bed was used as an initial translation transform, ensuring close-to-optimum initialization. Rigid, affine, and SyN registration were performed.

At the end of this step, a temporary full-block T_2_-weighted image was reconstructed to be used in the next registration step. To merge neighbouring FOVs, we applied a smooth weighted average in the overlap, giving more importance to voxels close to the isocentre of each FOV (so that voxels with a better signal are favoured).

#### Inter-Block registration

Inter-block registration consisted of registering non-overlapping reconstructed blocks, ensuring an accurate alignment along the cutting plane. It was the most delicate registration step of this whole pipeline: while image registration tools are widely available [46, 47], solutions for non-overlapping data are scarce and unsuited to our needs. Even if there was no overlap between the blocks after the cutting, we considered that, due to the continuity of brain structures, the cutting plane was an extremely thin overlap surface. By giving thickness to that surface, we enabled the use of regular 3D image registration techniques. We therefore developed a semi-automatic registration method based on a thick-surface representation of cutting planes. Figure 6 shows the outline of this method.

**Figure 6:**
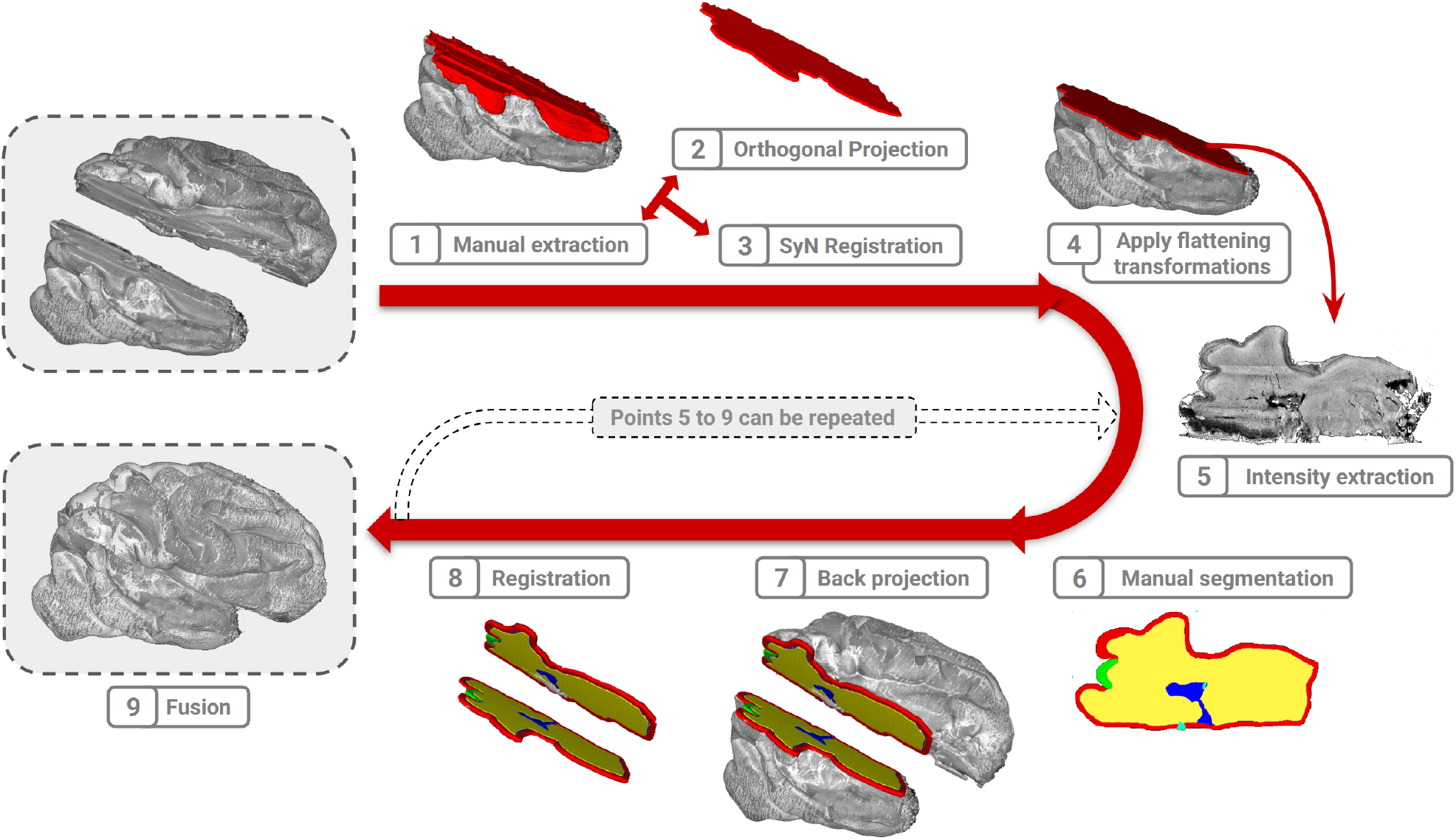
Schematic of the semi-automatic inter-block registration. Illustrated for sub-t27arh. Top left: start of the process—two blocks to be merged. Bottom left: reconstructed hemisphere. The numbering of steps corresponds to the main text under Materials and Methods – Inter-block registration.

1. As the brain is a malleable organ, the cutting surface, which was initially a plane, was no longer so at the time of imaging. We therefore sought to compensate for deformation after cutting. To that end, the cutting surface was first manually segmented in each block by using 3D Slicer [48].
2. To constrain the cutting surface to be flat, we first calculated the closest approximating plane by applying singular value decomposition (SVD) to the cutting surface segmentation performed in step 1. Voxels from the manual segmentation were orthogonally projected into the estimated plane, yielding a flat surface in 3D, with a thickness of 1 voxel (0.1 mm). Then, this projection was dilated to create a thick surface (target thickness of 1 mm).
3. Flattening transformations were generated by registering the manual segmentation (from step 1) onto the thick flat segmentation (from step 2) using SyN registration from ANTs [45] with increased regularization (cross-correlation cost function, gradient step = 0.5, update field variance = 6, total field variance = 1). By smoothing the update field variance, we sought to avoid strong local deformations, and by smoothing the total variance field, we sought to add “rigidity”.
4. The deformations calculated in the previous step were applied to the T_2_-weighted images of the blocks, to rectify their cutting surface into a planar geometry.
5. We then extracted the image intensities within the cutting plane from the rectified blocks, yielding a 2-dimensional (2D) image of each block along the cutting plane. To avoid the image being tainted by surface defects, we applied an offset to sample intensities at a depth of 0.5 mm (5 voxels) into the block.
6. On these 2D images, we performed a 2D manual segmentation of recognizable anatomical structures shared between both blocks. The exact labels used could vary, but commonly included: the cortical plate, ventricles, deep grey nuclei, vessels, specific gyri. The granularity of this labelling was refined as necessary, based on the final result, by repeating this and the subsequent steps.
7. The 2D manual segmentations were back projected into the 3D geometry of each block, by returning voxels to their initial coordinates in the rectified reconstructed block from step 4. The segmentation labels were then dilated to create a labelled thick surface.
8. Registration was performed between the two thick manual feature segmentations of adjacent blocks, using 3D isometric, affine, and SyN transformations (ANTs). The following parameters were used to perform the registration: cross-correlation cost function, gradient step = 0.5, update field variance = 6, total field variance = 1.
9. Finally, the transformations obtained at step 8 were applied to the T_2_-weighted images of both blocks, and intensities were merged by taking voxels with maximum intensity, yielding a temporary reconstruction of the adjacent blocks.

The reconstructed image was carefully inspected visually, and the manual segmentations were adjusted if necessary (redoing steps 6 to 9), until the resulting alignment of the blocks was deemed satisfactory.

#### Envelope correction

After inter-block registration had ensured local consistency along the cutting plane, the final step aimed to ensure the global consistency of the hemispheric geometry. Indeed, each block, having been acquired with a specific physical orientation in the scanner, could be slightly distorted along a different axis, resulting in an inconsistent geometry of the whole hemisphere. To correct for this inconsistency, rigid, affine, and SyN diffeomorphic registration (update field variance = 6, total field variance = 1) were computed between the binary mask of the reconstructed hemisphere and the mask of the 7 T reference image. Binary masks were used, rather than the T_2_-weighted images, to avoid introducing strong local deformations around deep structures whose geometry varies between the 7 T and 11.7 T images, e.g. the ventricles.

This final reconstruction space was rigidly reoriented into an anatomical AC-PC referential (Anterior Commissure—Posterior Commissure) centred on the AC, akin to an unscaled Talairach referential having standard RAS^+^ axes (Right, Anterior, Superior).

#### Jacobian maps

We concatenated all the deformation fields going from the final reconstruction space to the acquisition space of the T_2_-weighted reference, and for each voxel we calculated the natural logarithm of the determinant of the Jacobian matrix of that total deformation field ln(|J|). These maps quantify the logarithm of the local volume change (negative for expansion, positive for contraction) induced by the whole registration chain at each point of the reconstructed image.

### Image fusion

The registration protocol described above yielded a series of linear and non-linear spatial transformations that were used to resample all modalities into the final reconstruction referential, using trilinear interpolation. However, the number of valid measurements taken for each voxel of the reconstruction referential could vary: 1 in typical locations belonging to a single FOV, 2 in areas of FOV overlap, and occasionally 0 for voxels of the cutting plane that were corrupted by surface defects such as partial volume effect. Therefore, reconstructing a single whole-brain image for each modality required an image fusion strategy. A weighted-average strategy was used in areas of multiple measurements, where the weights were provided by a combination of geometric criteria and goodness-of-fit criteria when available. An inpainting strategy was used to fill small gaps along the cutting plane where no valid measurement was available. Log Jacobian maps used a simpler fusion strategy, selecting the measurement with the lowest absolute value.

#### Geometric weighting

For the fusion of all quantitative maps as well as anatomical images, a geometric weighting scheme was employed. Weights were based on the distance *d* of each voxel from the edge of the FOV, with the aim of emphasizing central regions that typically exhibit higher signal-to-noise ratio (SNR). To compute the weights *w*_geo_, the binary mask of each FOV was transformed into the final reconstruction space, and a Euclidean distance transform was applied replacing each voxel with its distance *d* to the edge of the FOV. The distance was subsequently passed through a sigmoid function (equation 1 with *d*_0_ = 9 mm, and *k* = 1.5) to produce a smooth transition, such that voxels within approximately 13 mm of the FOV edge experienced a gradual reduction in weight.

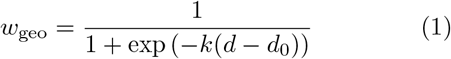

#### Goodness-of-fit weighting

In addition, the weights were further modulated based on a goodness-of-fit metric whenever available (i.e. for relaxometric maps). For each quantitative map, a residual map was computed by measuring the voxel-wise root-mean-square error (RMSE) between the measured signal curve and the fitted model. These residual maps, along with the quantitative maps, were transformed into the final space and spatially smoothed (*σ* = 0.2 mm). Given the smoothed residuals (*r*_1_, *r*_2_), the goodness-of-fit weight *w*_fit_ was defined as per equation 2.

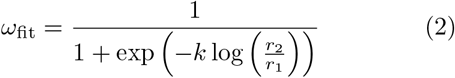

Here, the logarithm of the residual ratio ensures symmetry around zero, and the sigmoid function, with parameter *k* = 4, enables a smooth transition of weights favouring lower residuals, and hence more reliable estimates. In regions of overlap, voxel intensities were weighted with a combination of *w*_geo_ and *w*_fit_.

#### Inpainting of missing data

Along the cutting plane, some voxels of the reconstructed image space lacked a valid measurement from either block, because the voxels at the edge of the blocks were often corrupted by partial volume effect from non-brain tissue, typically a thin layer of PBS adhering to the surface of the tissue block. We constructed a mask of voxels with strong intensity variations by manually thresholding the Laplacian map so that most discontinuities were included, and limited the mask to 1.5 mm from the cutting plane. To enforce the continuity of the reconstructed data, we used biharmonic inpainting to replace voxels of this mask with plausible values interpolated from valid neighbouring voxels [49, 50] (scikit-image, RRID:SCR_021142).

### Data Records

The dataset [51] is available on the EBRAINS repository at https://doi.org/10.25493/X3K8-Y9W.

The dataset is organized according to the BIDS standard, version 1.10.0 [52, 53] (RRID:SCR_016124). An overview of the file structure is given in Figure 7, and the main files are described below. For any files not listed here, refer to the BIDS specification.

**Figure 7:**
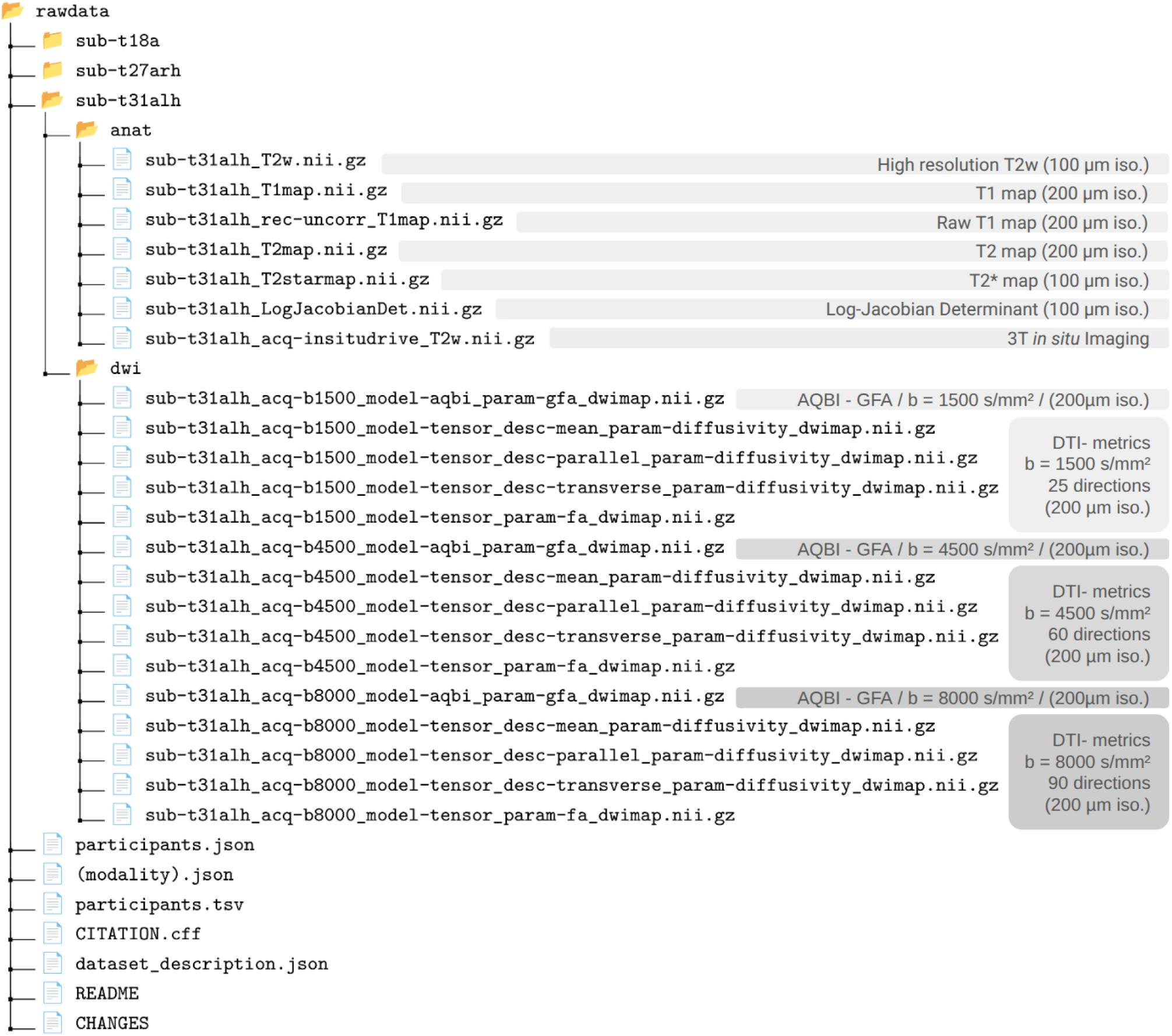
File organization of the dataset following the BIDS naming convention, as described in the Data Records section.

### General non-imaging data

- `README` and `CHANGES`: general information about the dataset and its usage; updates and complements to this data descriptor will be added to these files in future versions of the dataset.
- `dataset_description.json` and `CITATION.cff`: general metadata such as title, authors, and affiliations, funding, ethics, and links to the scripts and tools used to reconstruct the dataset.
- `participants.tsv` and `participants.json`: listing of the specimens and their general characteristics (similar to Table 1).

### Imaging data

As per the BIDS standard, images are stored in compressed NIfTI format, and filenames consist of a series of key-value pairs (described below) followed by a modality suffix: `T2w` for T_2_-weighted anatomical images, `T1map, T2map, T2starmap` for relaxometry maps. Data types not currently covered by the BIDS specification are also included: `dwimap` for maps derived from diffusion imaging (according to a work-in-progress nomenclature from BIDS Enhancement Proposal BEP016), and `LogJacobianDet` for the log-Jacobian determinant of the deformation field.

`sub-`is the identifier of the specimen according to the following naming convention: the letter t for *typical* (future releases may use other letters to identify brains with atypical neurodevelopment), + the rounded *age* in post-conceptional weeks, a *unicity letter* to ensure the unicity of the identifier, and optionally `lh` or `rh` corresponding to left hemisphere or right hemisphere, if only one hemisphere was imaged.

`acq-` disambiguates acquisitions of a given modality: `insitudrive` refers to the 3-tesla in situ acquisition; `b1500, b4500`, or `b8000` specify the b-value associated with the diffusion metrics.

`rec-` disambiguates multiple reconstructions of the same acquired data. Here `uncorr` was used to refer to the uncorrected T_1_ map (see Technical Validation—Relaxometry).

`model-`corresponds to the model used to reconstruct diffusion data: `aqbi` for Analytical Q-Ball Imaging, and dti for Diffusion Tensor Imaging.

`param-` is used to distinguish diffusion metrics: `diffusivity, fa` for Fractional Anisotropy, or gfa for Generalized Fractional Anisotropy.

`desc-` distinguishes types of diffusivity maps: `mean, parallel`, or `transverse` diffusivity.

### Technical Validation

Every step of the pipeline, from tissue preparation up to image registration and fusion, was implemented with the goal of optimizing the anatomical and microstructural fidelity of the final images to the original tissue samples. Best practices, quality checks, corrective measures, and limitations are described below.

#### Tissue integrity

Sample preparation and handling were conducted in accordance with the guidelines provided by the ISMRM Diffusion Study Group for preclinical diffusion MRI [54], as closely as possible. Despite adherence to these recommendations and the implementation of multiple quality control steps, the tissue samples were variably affected by unavoidable factors such as the conditions surrounding death, post-mortem intervals (PMI), and handling-related stress. As highlighted in the ISMRM guidelines, minimizing the PMI is crucial to preserving tissue integrity; however, in the context of fetal death, clinical circumstances cannot be anticipated nor controlled, even with the best efforts of the medical staff. These factors are inherent to any ex vivo study on human fetal tissue and could have led to some deterioration of the tissue microstructure, which may limit the generalizability of findings in areas of altered tissue. These deteriorations are particularly visible in the diffusion metrics, barely visible in relaxometry maps and invisible in the T_2_-weighted images.

Fetal tissue, being mostly unmyelinated, is particularly fragile and prone to tearing under its own weight. Keeping the arachnoid matter attached to the hemispheres helps to preserve mechanical integrity of the tissue—it was unfortunately removed from the three specimens of this first data release. Despite the most careful handling, all hemispheres exhibited localized macroscopic damage: the affected regions are listed in `participants.tsv` under the `deteriorations`column.

### Acquisition & parametric mapping

#### Preliminary acquisitions

During gestation, the fetal brain undergoes drastic changes in composition, volume and structure. Therefore, to ensure that the imaging sequences provide optimized signal and contrast, we performed a preliminary acquisition of each specimen to estimate whole-brain individual relaxation times at 11.7 T. We assessed T_1_ and T_2_ relaxation times using 2D inversion recovery and variable echo time EPI sequences on a few slices, with millimetric spatial resolution. We fixed the sequence parameters described in the Methods using the first specimen, and none of the subsequent ones showed strongly deviant values necessitating a change of sequence parameters. Using this technique before and after gadolinium doping, we also confirmed that relaxation times were greatly reduced by doping: T_1_ by about 90%, T_2_ by about 40%.

The choice of b-values for diffusion imaging was also based on a preliminary acquisition: we ran a succession of PGSE sequences sampling the b-value every 500 to 1000 s/mm^2^ from 500 to 8000 s/mm^2^. We then plotted the log(*S*/*S*_0_) curve as a function of b-value for different types of tissues (see Figure 8). For the previously described protocol, we chose b-values that captured the initial linear diffusion regime—b = 0 s/mm^2^ and b = 1500 s/mm^2^—as well as the non-linear regime—b = 4500 s/mm^2^ and b = 8000 s/mm^2^. This choice will enable the use of advanced microstructure models such as Neurite Orientation Distribution and Density Index (NODDI) [55], as well as high angular resolution tractography thanks to the high b-value.

**Figure 8:**
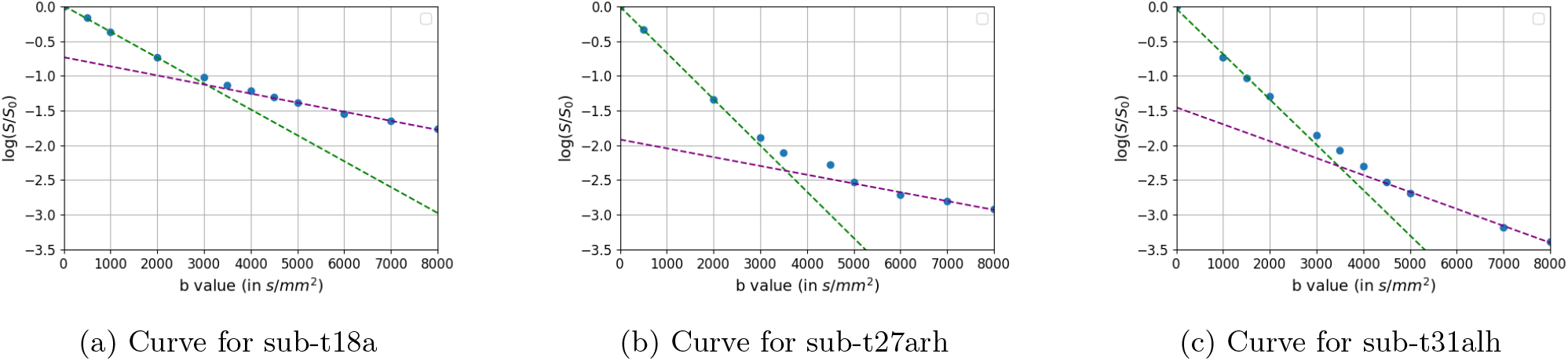
Evolution of the diffusion signal *S* with the b-value, measured during preliminary acquisitions. Green: linear regression on the low b-values (initial linear diffusion regime). Purple: linear regression on the high b-values points (asymptotic regime). The data are extracted from the intermediate zone for sub-t18a and sub-t27arh, and from the white matter for sub-t31alh. It should be noted that the difference in the appearance of the curves may be related to different water content in the tissues.

#### Scanner limitations

In this study, we pushed the technical capabilities of the preclinical MRI scanner by acquiring FOVs larger than originally intended by the manufacturer. Notably, the quality of the gradients is guaranteed only within a 3 cm diameter sphere around the isocentre (diameter of the sphere volume, where the manufacturer specifies 10% maximum deviation from the ideal gradient). Accordingly, strong geometric distortions, on the order of several millimetres, were observed at the FOV edges (typical FOV length was 6 cm). Gradient non-linearity correction was able to completely unwarp the images, as evidenced by the success of inter-FOV registration, which displayed sub-voxel accuracy, even when only using an affine transformation.

The radiofrequency (RF) coil was also used at its limits. As a result, transmission and reception efficacy could vary greatly between voxels located at the centre of the FOV and those at the periphery (FOV length is typically 6 cm). Reception bias was inherently compensated for by quantitative relaxometry and diffusion models, and by an explicit bias correction step (N4) for the T_2_-weighted anatomical image. On the other hand, transmission inhomogeneities more fundamentally affect signal production: this and other phenomena can adversely affect the quality of the reconstructions. The rest of this section will list the issues encountered with each modality and describe the corrective measures taken.

#### T_1_ relaxometry

To compensate for RF transmission inhomogeneities, an explicit correction using a 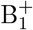 map was implemented as described in the Methods. However, the T_1_ maps still exhibited inhomogeneities across the tissue, mostly appearing as a vertical gradient with very low T_1_ values toward the top of each FOV. The cause of this gradient of T_1_ could not be fully elucidated. It was not due to incomplete correction of a strong 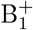 inhomogeneity, as the regions of low T_1_ did not spatially correspond with the pattern of extreme 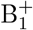 values (see Figure 9), even when using a different 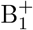 -mapping method (the double angle method)[56]. One hypothesis is that these T_1_ measurement variations result from a vertical gradient of both tissue hydration and Gadolinium concentration, induced by a combination of gravity and buoyancy forces acting onto the tissue.

**Figure 9:**
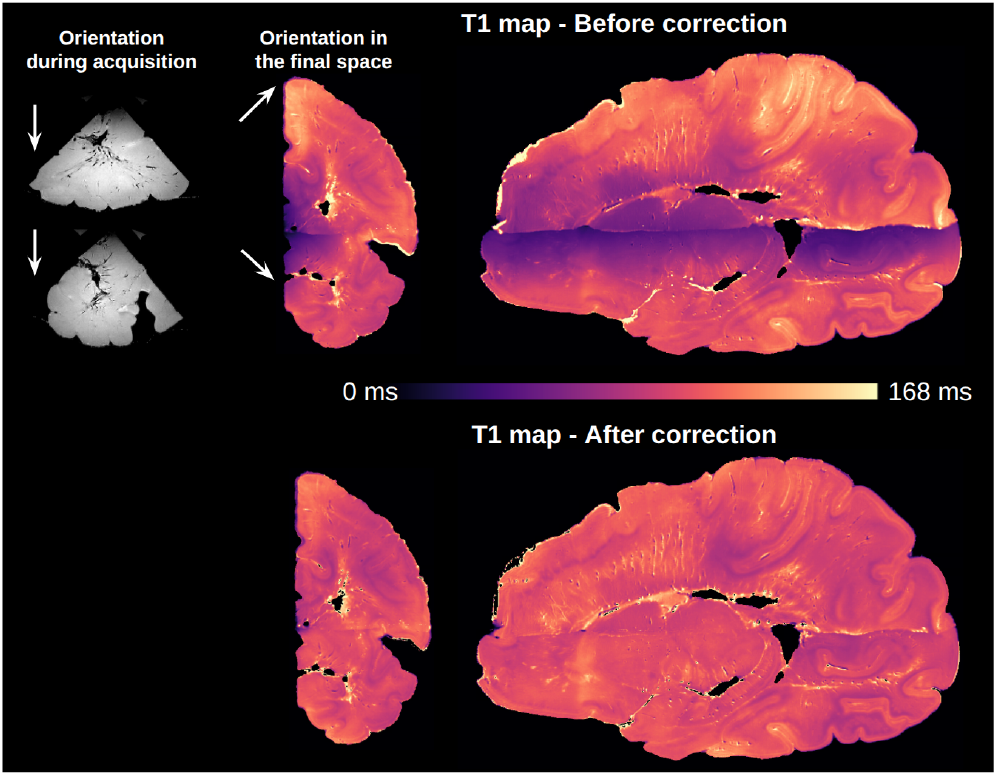
Bias correction of T_1_ maps, illustrated with sub-t31a. Upper left: orientation of the blocks in the initial space. Upper middle and right: coronal and sagittal views of the uncorrected T_1_ map in the final space. White arrows indicate the vertical direction (gravity) for each block. Bottom: coronal and sagittal views of the bias-corrected T_1_ map.

As this gradient of T_1_ had no plausible physiological origin, we applied a multiplicative bias correction (see Methods) with satisfactory results, allowing to recover similar values and contrast in the low-T_1_ regions compared to the rest of the tissue (see Figure 9).

#### T_2_ relaxometry

The T_2_-mapping sequence, which uses a train of refocusing pulses to achieve multiple echo times, is prone to artefacts due to transmit field inhomogeneity. Indeed, we observed artefacts that increased with successive echoes and that corresponded to the pattern of the B_1_ map. These artefacts introduced biases in the estimation of T_2_ values. Unfortunately, correcting them, if at all possible, would have required advanced signal modelling methods such as extended phase graphs, which were not possible within the time frame of this work. Therefore, caution should be exercised when interpreting the published T_2_ maps reconstructed by a simple monoexponential fitting, as they remain affected by a residual bias.

#### 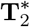 relaxometry

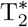 relaxometry is robust to transmission inhomogeneities, as the multiple echoes are acquired after a single excitation pulse. This modality is sensitive to inhomogeneities of the static magnetic field B_0_, particularly those created by air bubbles and blood byproducts, but this was mitigated by the use of a small voxel size (100 µm) that greatly limits signal loss due to intra-voxel dephasing.

#### Diffusion

We found the diffusion-unweighted (b = 0 s/mm^2^) images to be affected by a systematic signal drop, appearing at a consistent spatial location relative to the scanner, regardless of the specimen or the position of the tissue (illustrated in Figure 10). The diffusion-weighted images were not affected by this signal drop, leading to paradoxical situations where *S*_b=0_ *> S*_b=0_. The origin of that signal drop remained unresolved, but the fact that its location was scanner-based rather than tissue-based indicated an issue with the acquisition. The multiplicative bias-correction strategy described in the Methods proved effective on the b=0 images and allowed us to successfully fit the DTI and AQBI diffusion models, with a satisfactory homogeneity of the resulting metrics (see Figure 10).

**Figure 10:**
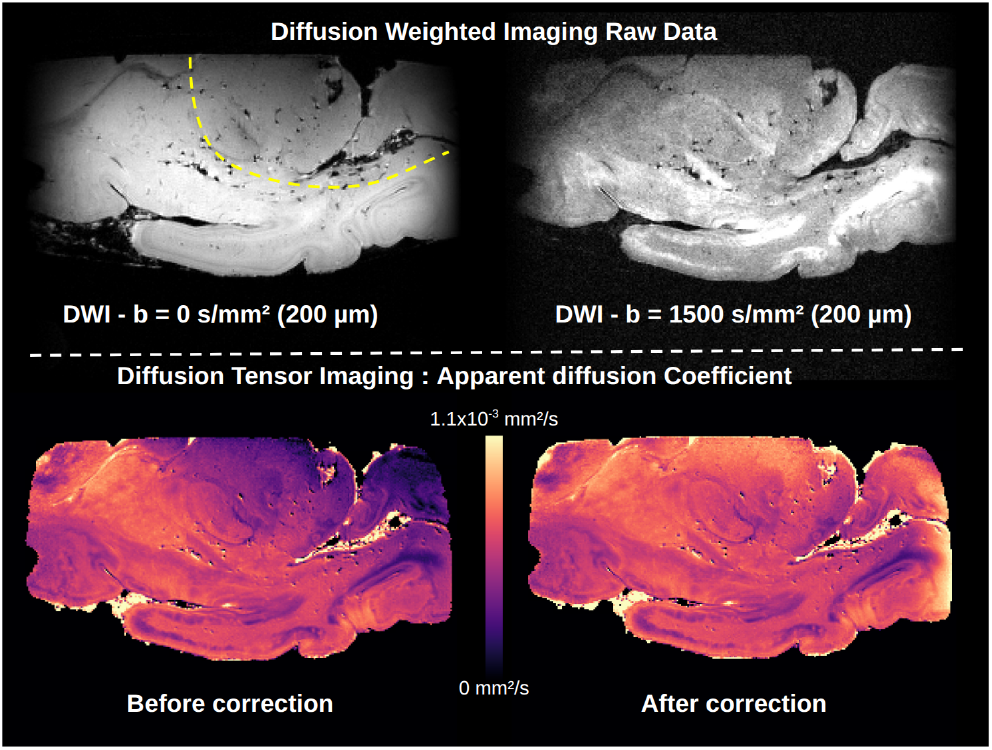
Bias correction of the diffusion-unweighted (b = 0) image, illustrated on the inferior block of sub-t31alh. Top left: raw data at b = 0 s/mm^2^ (oblique view) with the hypointense region delimited by the yellow dotted line. Top right: raw data at b = 1500 s/mm^2^ showing no inhomogeneity. Bottom left: apparent diffusion coefficient before correction. Bottom right: apparent diffusion coefficient after correction. DWI = diffusion weighted imaging.

For each diffusion-weighted image of each FOV, a visual quality check was performed to assess the effectiveness of motion and eddy current artefact correction. The consistency of diffusivity values was verified by calculating the mean diffusivity within liquid-filled voxels, located either in the ventricles or in the retention gutter of the container. A value of 1.98 *×* 10^−3^ mm^2^/s was obtained, which corresponds closely to the known diffusivity of water at 20°C—approximately the ambient temperature during image acquisition.

Despite our rigorous analysis pipeline and corrections of residual artefacts, we identified abnormal patterns in some regions of the reconstructed maps, particularly in areas of subcortical white matter, as illustrated in Figure 11. These patterns may suggest alterations to the microstructural integrity of the tissue, related to either peri-mortem pathological processes or post-mortem microstructural alterations. In particular, very low diffusivities and high fractional anisotropy may result from such abnormal artefactual processes. Diffusion data from those regions should not be included in analyses seeking to describe the non-pathological brain.

**Figure 11:**
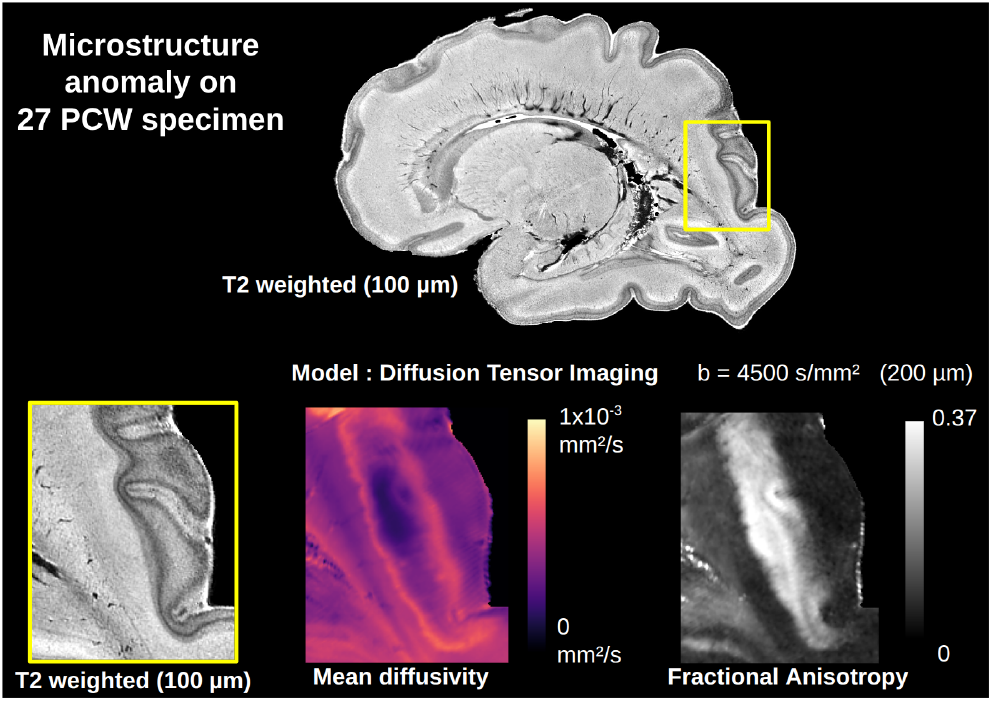
Tissues anomalies seen on diffusion images. (as visible on sub-t27arh). Top: whole-hemisphere T_2_-weighted sagittal slice. Bottom row: zoom on a subcortical region. The high-resolution T_2_-weighted image shows no particular pattern but diffusion metrics at b = 4500 s/mm^2^ show FA and mean diffusivity anomalies in the subcortical region.

### Registration & Fusion

Throughout the whole acquisition and reconstruction pipeline, each step was implemented with the aim of avoiding unnecessary deformation of the images. That effort began at the stage of tissue preparation and conditioning: fetal brain tissue being extremely soft, the use of custom-fitted 3D-printed containers was essential to avoid introducing deformations that would later need to be corrected by image registration. For registration steps, spatial masks were used whenever possible to spatially constrain the evaluation of the cost function to regions where geometric corrections were required. Furthermore, the choice of the T_2_-weighted modality as a reference hub for all other modalities enabled the use of monomodal registration throughout, which is typically more robust and accurate than cross-modal registration, thereby limiting the amount of noise introduced into the deformation fields.

Objective evaluation of image registration is notoriously difficult, particularly in such cases where no ground truth is available. Here, we calculated the log Jacobian determinant of the total deformation field between the acquired FOVs and the reconstructed image, providing a quantitative characterization of the deformation induced by the registration pipeline. On these maps, smooth low-amplitude variations in the bulk of the tissue are a good indication that no strong local deformations were introduced in those areas. On the other hand, higher values with steeper variations are observed where deformations were implemented to match the geometry of the tissue blocks: typically along the cutting planes, and close to structurally unconstrained parts of the tissue (e.g. the ventricles). These maps could be used to inform downstream analyses, either by accounting for the induced deformations or by deciding to exclude strongly warped regions of the tissue.

### Usage Notes

The mesoscopic isotropic resolution, the multimodality, and the whole-hemispheric coverage of this unique dataset of post-mortem developing brains make it an ideal candidate to serve as a reference dataset for studying fetal neurodevelopment. It may serve as a linking hub to provide a common coordinate framework to a wide range of studies, ranging from microscopic-scale data, such as immunohistochemical images and, within the limits of comparability, to neuroimaging data from the entire brain population, such as in utero image cohorts. The integration of this dataset into the EBRAINS interactive atlas viewer will facilitate such use cases.

Beyond educational purposes, this dataset also offers a unique opportunity to explore neurodevelopmental processes at the mesoscopic scale through a multimodal approach. In addition to anatomical T_2_-weighted images with unprecedented detail, the combination of relaxometry 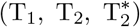 and diffusion tensor imaging (DTI) provides rich, versatile, quantitative proxies of tissue microstructure. These modalities, when analysed in concert, offer strong descriptive potential for characterizing the structural and connectivity maturation of the brain during the second half of gestation (see Figure 2). By analysing the variations of these proxies across brains at different developmental stages, neuroscientists can identify and quantify global and local dynamic patterns that are otherwise difficult to capture.

Moreover, the mesoscopic resolution opens access to the study of the cerebral vasculature. Features related to vessel morphology, density, and spatial distribution—particularly visible on 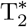 and DTI contrasts—can be analysed in detail. This offers new perspectives on the interaction between vascular development and brain maturation.

The T_2_-weighted images and 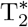 maps showed consistently good signal quality. For other modalities, we encourage users of the data to carefully consider the limitations outlined in the Technical Validation. To summarize, users should be mindful of the regional corrections that were applied to the T_1_ maps and b = 0 s/mm^2^ images. Our T_2_ maps are known to suffer from spatial inhomogeneity that may limit their interpretation. Moreover, particular attention should be given to the potential alterations in microstructure due to peri-mortem and post-mortem tissue degradation, as suggested by the diffusion metrics: we recommend using the mean diffusivity to identify regions affected by those alterations. Additionally, values in voxels located along the cutting planes are known to be less reliable than in other regions: they can result from inpainting or be affected by strong local deformations. The log-Jacobian maps can be used to identify strongly warped regions and treat them appropriately.

Future data releases will enrich this dataset in at least four directions: i. additional brains of different developmental ages will be added, which will improve the coverage of the developmental timeline from mid-gestation all the way to term; ii. multi-shell diffusion microstructure models such as NODDI will be fitted, and a pipeline will be developed to reconstruct orientation distribution functions (ODFs) from the blockwise diffusion data, enabling the study of fibre architecture through tractography; iii. co-registered histological images acquired on the same brains will be provided as histological ground truth; iv. a segmentation of brain structures such as transitory compartments (ventricular zone, intermediate zone, etc.) will be performed and released, turning this multimodal MRI dataset into a full-featured developmental atlas.

## Data availability

The dataset described by this article [51] is publicly distributed on the EBRAINS repository at https://doi.org/10.25493/X3K8-Y9W.

## Code availability

Scripts used for all data processing are available in the accompanying repository: https://github.com/neurospin/phcp-reco, which includes detailed instructions for use. The semi-automatic inter-block registration process requires frequent manual inputs, and is therefore provided as a collection of modular processing units rather than a fixed pipeline. The quantitative maps were reconstructed using algorithms provided by the Ginkgo toolbox: https://framagit.org/cpoupon/gkg.

## Acknowledgements

We would like to thank the medical staff, MR technicians and members of the fetal pathology laboratory at Robert-Debré Hospital for their hard work and efficiency that were critically useful to this project. We thank Ivy Uszynski and Simon Legeay for their technical advices, as well as Lina Abdallah-Khodja for her critical feedback on the data, and Naz Karadag from the EBRAINS curation team for the careful review of this data descriptor. We thank Jérémy Bernard for designing the external container components and for his assistance with 3D printing, as well as Benjamin Lehmann for printing many of the custom-made devices. This work was funded by a grant from the French *Agence Nationale de la Recherche* (ANR) for the p-HCP project (ANR-21-CE37-0029); and benefited from a government grant under the France 2030 program, under the reference ANR-23-IAHU-0010.

## Author contributions

Conceptualization: C.P., H.A.-B., L.H.-P., J.D., Y.L. Methodology: L.A., C.P., Y.L. Resources: S.K.-S., H.A.-B. Investigation: L.A., S.K.-S., Y.L., M.A. Software: L.A., C.P., Y.L. Formal analysis: L.A. Validation: L.H.-P., Y.L., J.D. Data curation: Y.L., L.A. Visualization: L.A., L.H.-P., Y.L. Writing – original draft: L.A. Writing – review & editing: all authors. Supervision: L.H.-P., Y.L., S.K.-S., M.A. Project administration: H.A.-B., L.H.-P. Funding acquisition: H.A.-B., C.P., L.H.-P., J.D.

## Competing interests

The authors declare no competing interests.

## Footnotes

1 Gestational age can be approximated by adding 2 weeks to the post-conceptional age, corresponding to the time elapsed since the last menstrual period.

## Notes

https://doi.org/10.25493/X3K8-Y9W

https://github.com/neurospin/phcp-reco

### Competing Interest Statement

The authors have declared no competing interest.

### Summary of Updates

Links to the dataset have been updated with the dataset DOI, and with the dataset version assigned by EBRAINS.

